# Stringent structural plasticity of dendritic spines revealed by two-photon glutamate uncaging in adult mouse neocortex *in vivo*

**DOI:** 10.1101/577742

**Authors:** Jun Noguchi, Akira Nagaoka, Tatsuya Hayama, Hasan Ucar, Sho Yagishita, Noriko Takahashi, Haruo Kasai

## Abstract

Two-photon uncaging of glutamate is widely utilized to characterize structural plasticity in brain slice preparations *in vitro*. In this study, we investigated spine plasticity by using, for the first time, glutamate uncaging in the neocortex of adult mice *in vivo*. Spine enlargement was successfully induced in a smaller fraction of spines in the neocortex (22%) than in young hippocampal slices (95%), even under a low magnesium condition. Once induced, the time course and mean amplitudes of long-term enlargement were the same (81%) as those *in vitro*. However, low-frequency (1–2 Hz) glutamate uncaging caused spine shrinkage in a similar fraction (34%) of spines as *in vitro*, but spread to the neighboring spines less frequently than *in vitro*. Thus, we found that structural plasticity can occur similarly in the adult neocortex *in vivo* as in the hippocampus *in vitro*, although it happens stringently in a smaller subset of spines.

## Introduction

Most excitatory synapses in the brain form on dendritic spines. The volume of dendritic spines is tightly correlated with the functional expression of glutamate receptors in the young hippocampus *in vitro* (Matsuzaki et al., 2001; Smith et al., 2003; Beique et al., 2006; Asrican et al., 2007; Holbro et al., 2009; Zito et al., 2009) and in the adult mouse neocortex *in vivo* (Noguchi et al., 2011). Spine volume changes have been associated with long-term potentiation and depression of synapses in hippocampal preparations (Zhou et al., 2004; Kopec et al., 2007). Such volume changes eventually cause the generation and elimination of spines (Yasumatsu et al., 2008; Bhatt et al., 2009; Holtmaat et al., 2009; Xu et al., 2009; Kasai et al., 2010; Hayashi-Takagi et al., 2015). Impaired structural plasticity induces pathological states of neuronal circuits (Fiala et al., 2002; Kasai et al., 2010; Forrest et al., 2018).

Two-photon uncaging of caged glutamate compounds (Matsuzaki et al., 2001) is the only available method that reliably stimulates single spines. It is widely used to characterize spine structural plasticity *in vitro*. Spine enlargement is most robustly induced by uncaging caged glutamate in the absence of external magnesium (Mg^2+^) so that *N*-methyl-D-aspartic acid (NMDA) receptors are maximally activated (Matsuzaki et al., 2004; Noguchi et al., 2005; Harvey et al., 2007; Honkura et al., 2008; Lee et al., 2009; Govindarajan et al., 2011; Bosch et al., 2014). Spine shrinkage can be induced by low-frequency uncaging (Hayama et al., 2013; Oh et al., 2013; Noguchi et al., 2016). However, assessing spine plasticity with two-photon uncaging has never been characterized *in vivo* because of difficulties in uncaging *in vivo*. The characteristics of structural plasticity is unknown in the adult mouse neocortex *in vivo*.

We previously established a glutamate uncaging method *in vivo* in which a caged glutamate compound is applied on the surface of the brain. This method allows the compound to spread into the superficial extracellular space of the neocortex for free diffusion (Noguchi et al., 2011). We now extend our study to focus on the structural plasticity of dendritic spines *in vivo*.

## Results and Discussion

### Spine enlargement *in vivo*

Two-photon uncaging of the caged glutamate compound was applied to single spines of tuft dendrites of layer 5/6 pyramidal neurons in the visual cortex of adult mice *in vivo* (Noguchi et al., 2011). We used the green fluorescent protein (GFP)-expressing M mouse line or the yellow fluorescent protein (YFP)-expressing H mouse line in which a subset of layer 5/6 pyramidal neurons was selectively labelled. Mice were anesthetized with urethane and xylazine and placed under a microscope objective lens using an imaging chamber that was firmly attached on the skull of the mouse (*Figure 1A*). To activate NMDA receptors effectively, the recording chamber was superfused with artificial cerebrospinal fluid containing no magnesium (Mg) ions. Caged glutamate was thereafter superfused (*Figure 1A and Figure 1–figure supplement 1A*). Spine head volume (V_H_) fluctuations before uncaging were quantified as coefficients of variation (CVs) (*Figure 1–figure supplement 1B*). The CV of *in vivo* neocortex spines (15% ± 16% [mean ± standard deviation (SD)]; 227 spines) was not larger than that of hippocampal slices (21%)(Matsuzaki et al., 2004), which ensured the stability of our recording conditions.

**Figure 1.**
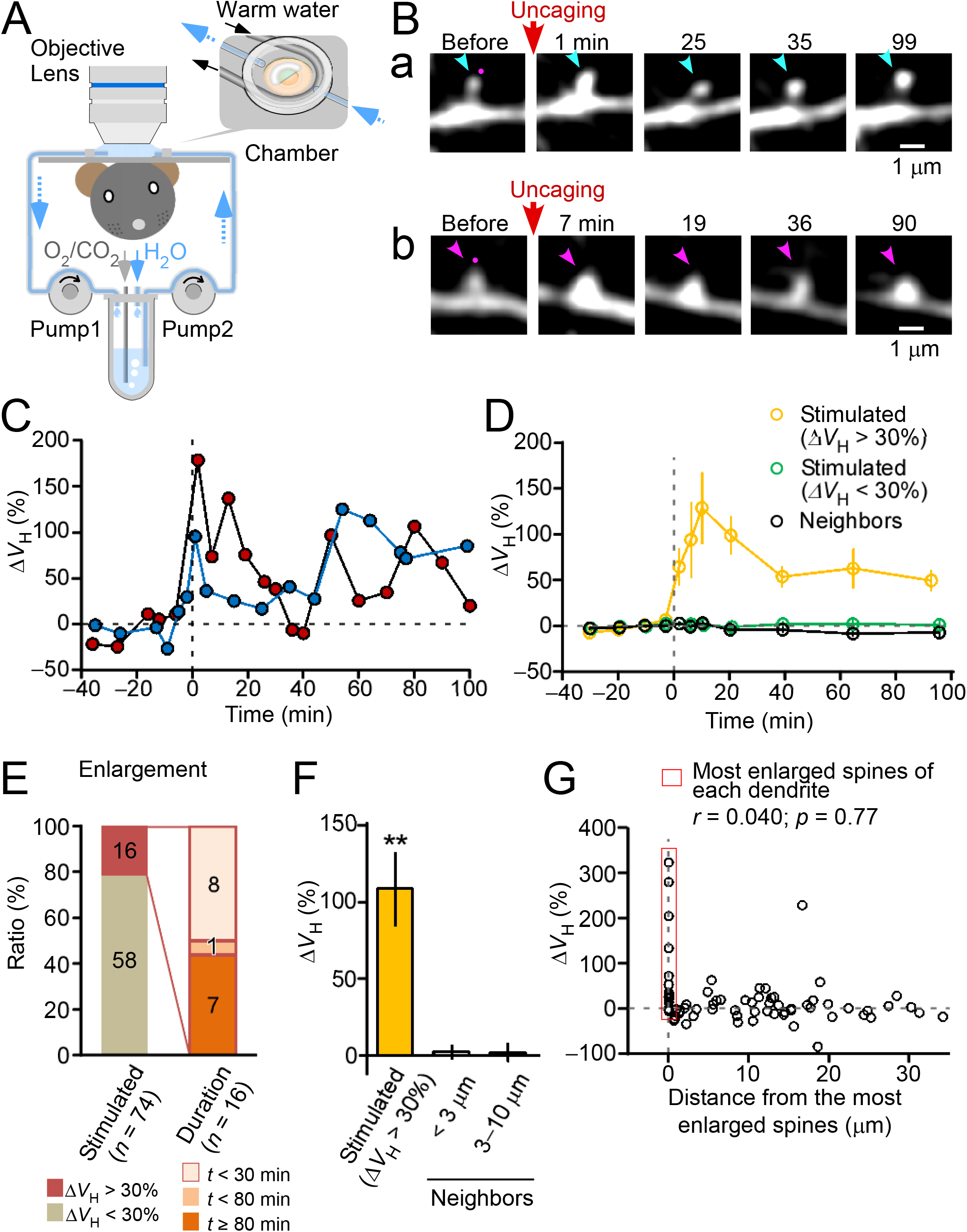
Induction of spine enlargement in the visual cortex *in vivo*. (**A**) Schematic drawing of the experiment. Transgenic mice that expressed green fluorescent protein (GFP) or yellow fluorescent protein (YFP) in neocortex layer 5/6 pyramidal neurons were urethane-anesthetized and placed under an objective lens using a metal chamber. Skull and dura over the neocortex V1 area were carefully removed, and a half-moon-shaped coverslip was placed on the brain surface. A perfusion solution containing caged glutamate and 10 μM tetrodotoxin (TTX), but no magnesium (Mg^2+^) ions, was steadily circulated using peristaltic pumps. After diffusing the caged glutamate into the brain parenchyma, caged glutamate was photolyzed at the tip of dendritic spines by using two-photon uncaging at the wavelength of 720 nm. Dendrite images were obtained by another laser light path (see the “Materials and Methods” section for details). (**B**) Time-lapse images of the stimulated spines. Several spines (average, 4.5 spines) in each dendrite have been stimulated with repetitive two-photon glutamate uncaging. The red dots show the position of the uncaging. The blue and red arrowheads show the stimulated spines. (**C**) Time courses in the increase in the volume of spine a (blue) and spine b (red) in panel B. (**D**) The averaged time courses of the spine-head volume increment are plotted for spines with >30% enlargement (orange circles), spines with <30% enlargement (green circles), and unstimulated neighbor spines (black circles) (*n* = 16, *n* = 68, and *n* = 92 spines for enlarged spines, unenlarged spines, and neighbors, respectively). (**E**) Ratio of the change in head volume (ΔV_H_) >30% in the enlarged spines and the longevity of the enlargement. The left stacked bar presents the ratio of the enlarged spines (22%) to the remaining spines (78%) of all (20) dendrites. The right stacked bar presents the distribution of the enlargement duration. The numbers in the histograms indicate the number of spines. (**F**) The average increases in the spine volumes (109% ± 24%) in 16 stimulated spines and in neighboring spines located <3 μm (2.2% ± 4.6%; 12 spines) and at 3–10 μm (2.0% ± 6.0%; 8 spines). \*\**p* < 0.01, based on Wilcoxon signed-rank test against zero. The error bars represent the standard error of the mean (SEM). (**G**) Scatter plots of the average spine enlargement (10–30 min after stimulation [i.e., ΔV_H_]) of the stimulated spines against the distance between the most shrunken spine of each dendrite and other stimulated spines. Pearson’s product-moment correlation coefficient was calculated for the scatter plots.

Enlargement of the spines could be induced by two-photon glutamate uncaging repeated 60 times at 1 Hz adjacent to the spine heads (*Figures 1B and 1C*). Volume changes varied among individual spines; however, the averaged time course showed a transient increment phase, followed by a stable plateau phase (*Figure 1D*). For spines showing >30% enlargement, the peak enlargement (10–30 min) and sustained phase of enlargement (>60 min) were 109% ± 24% (the mean ± the standard error of the mean: 16 spines/10 dendrites/10 mice) and 50% ± 12%, respectively. These values were similar to those of CA1 pyramidal neurons *in vitro* (Matsuzaki et al., 2004). Enlargement lasting more than 30 min occurred in 8 of 16 enlarged spines (*Figure 1E*) and was confined to the stimulated spines (*Figures 1D and 1F*).

Enlargement was recorded only in a small fraction of spines (22% of 74 spines/20 dendrites/18 mice; *Figure 1E*), compared with the fraction in the hippocampus *in vitro* (approximately 95%) (Matsuzaki et al., 2004). In spines without enlargement (ΔV_H_ <30%), the average enlargement was negligible (−0.6% ± 2.5%) (*Figure 1D*). The stringency in spine enlargement was not because of technical reasons; the enlargement was induced mostly in one spine (0–4 spines; average, 0.8 spine) among several spines (1–7 spines; average, 3.7 spines) that were simultaneously stimulated. This conclusion was quantitatively supported by the fact that the amplitude of the enlargement of stimulated spines was uncorrelated with the distance of the spine from a spine showing significant enlargement (*Figure 1G*). In these studies, we selected small spines (*Figure 1–figure supplement 2A*) in which enlargement could be induced in the most pronounced manner. The enlargement was uncorrelated with spine neck length, depth, and mouse age (*Figures 1–figure supplement 2B-D*).

### Spine shrinkage *in vivo*

We used a solution containing a physiological concentration (1 mM) of Mg^2+^ to study spine shrinkage (Noguchi et al., 2016). Several spines on a dendrite were stimulated by low-frequency two-photon glutamate uncaging (2.8 spines/dendrite, 1–2 Hz, 5–15 min) (*Figure 2A*). Stimulated spines showed a large volume reduction (*Figures 2A and 2B*, spine “S1”). The spine volume gradually reduced *in vitro* (Hayama et al., 2013; Noguchi et al., 2016) (*Figure 2C*). We found that 34% of the stimulated spines had shrunk (–ΔV_H_ >30%, 15 of 44 spines/18 dendrites/8 mice) and that the mean amplitude at 20–50 min was 19% ± 4% (*n* = 44), which was similar to the findings of the young hippocampus *in vitro* (23% ± 7%, *n* = 8) (Noguchi et al., 2011). The shrinkage was long-lived (>80 min) in most (73%) spines (*Figure 2D*). Shrinkage was absent when we added the NMDA receptor antagonist APV in the perfusion solution (*Figure 2C; Figure 2–figure supplement 1A*).

**Figure 2.**
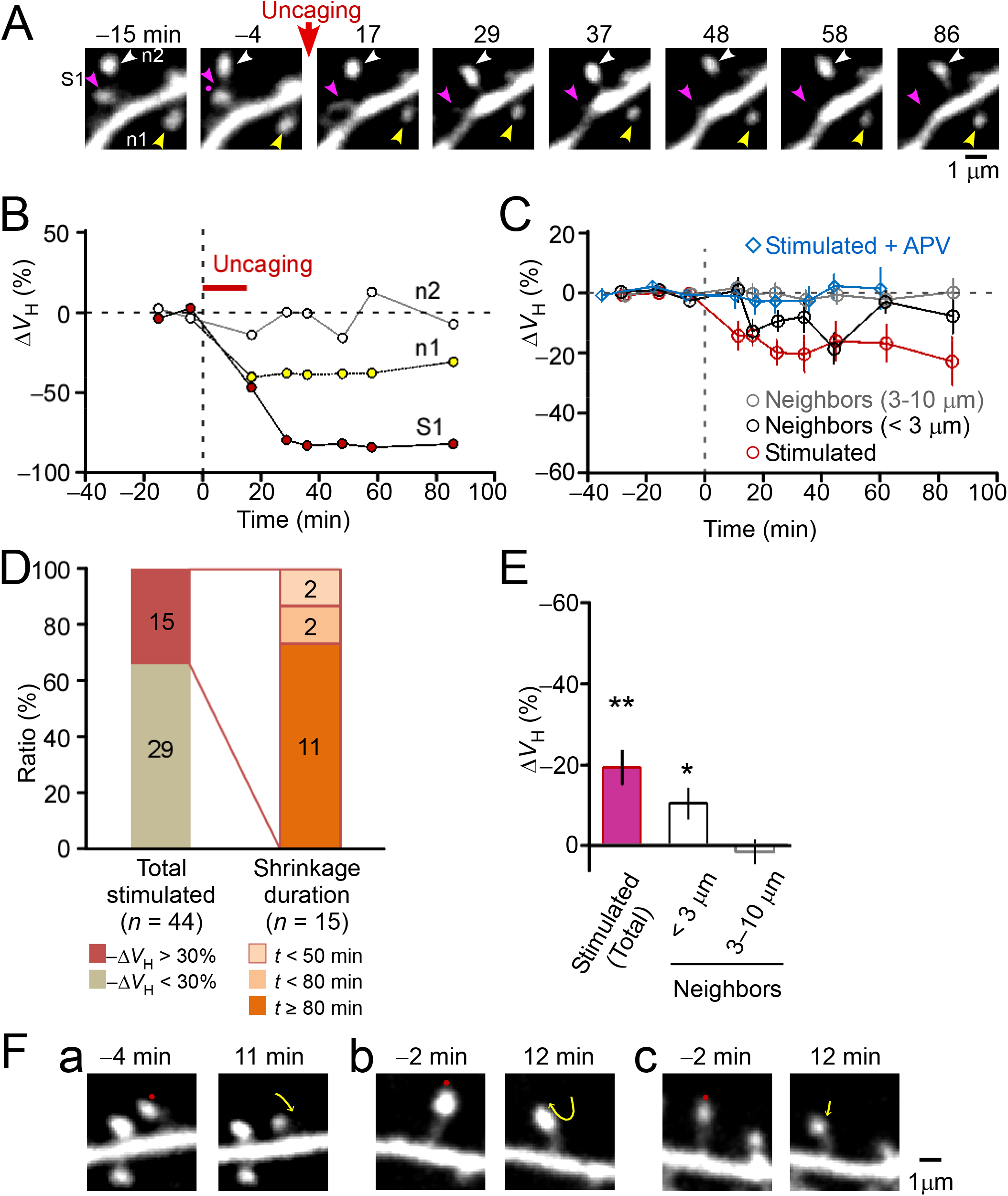
Induction of spine shrinkage *in vivo*. (**A**) Representative images of spine shrinkage. We stimulated, on average, 2.8 spines in a dendrite using the method used for the enlargement experiments. However, the perfusion solution contained 1 mM magnesium (Mg^2+^). Spines (S1, red arrowheads) stimulated with low-frequency two-photon glutamate uncaging (1 Hz, 15 min) show significant shrinkage. A neighboring spine is also shrunken (n1, yellow arrowheads) but another neighboring spine is not shrunken (n2, white arrowheads). The uncaging point is indicated by a small red dot. (**B**) Time-courses of the spine head volumes in panel A. The red, yellow, and white circles indicate the traces of spines S1, n1 and n2, respectively. (**C**) Average time courses for the stimulated spines without (red circles) or with the NMDA receptor antagonist APV (blue diamonds). Forty-four spines were not exposed to APV and 12 spines were exposed to APV. Average time courses of the neighbors located <3 μm (black circle) or 3–10 μm (gray circle) from the stimulated spines are also plotted (58 spines for <3 μm and 64 spines for 3–10 μm). (**D**) The ratio of the shrinkage in the volume head (-ΔV_H_) >30% in the stimulated spines and the longevity of the shrinkage. The left stacked bar chart presents the ratio of shrunken spines to the remaining spines of all (16) dendrites. The right chart presents the distribution of the shrinkage duration. The numbers in the bars indicate the number of spines. (**E**) The average amplitude of shrinkage of the stimulated spines (−19.1% ± 4.3%, 44 spines) and the neighboring spines <3 μm (−10.2% ± 3.8%, 58 spines) or 3–10 μm (1.8% ± 3.5%, 64 spines) from the stimulated spines. \**p* < 0.05 and \*\**p* < 0.01, based on Wilcoxon signed-rank test against zero. The error bars represent the standard error of the mean (SEM). (**F**) Three representative images of spine retraction after the low-frequency stimulation. The red dots and yellow arrows represent the uncaging points and direction of the retraction, respectively.

Spine shrinkage spread to neighboring spines, which also occurred in hippocampal slice culture samples (Hayama et al., 2013; Noguchi et al., 2016). We calculated the average spine volume of the stimulated spines and the neighboring spines at 20–50 min from the onset of stimulation (*Figure 2E*). We found that the diffusion of spine shrinkage was only significant in spines next to the stimulated spines (<3 μm). Only 12% of spines within 3 μm of a stimulated spine showed shrinkage (–ΔV_Stimulated_ >30%; *Figure 2–figure supplement 1B*). Thus, the spread of spine shrinkage was more stringent *in vivo* than *in vitro* in which shrinkage spread to 71% of spines within 3 μm from a stimulated spine and to 38% of spines within 7 μm (Hayama et al., 2013; Noguchi et al., 2016).

We found that the prestimulation spine volume showed a weak but insignificant correlation with spine shrinkage (*Figure 2–figure supplement 1C*). Spine retraction occurred during spine shrinkage (*Figure 2F*) (Hayama et al., 2013); however, spine shrinkage was insignificantly correlated with retraction (ΔSpine length; *Figure 2–figure supplement 1D*). We did not observe any interspine distance dependency in the induction of spine shrinkage (*Figure 2–figure supplement 2A*). Spine shrinkage was also insignificantly correlated with the initial spine neck length, dendritic depth, and age of mice (*Figure 2–figure supplement 2B–D*).

### Stringent structural plasticity of dendritic spines in the neocortex *in vivo*

We found that two-photon uncaging could induce prominent plasticity of spine structures in the adult neocortex *in vivo* that was similar to that of the hippocampus *in vitro*. The major difference was the low success rate of spine enlargement *in vivo* (22% vs. 95% *in vitro*), which was not caused by technical factors (*Figure 1G*). The success rate in inducing shrinkage was similar to that of the hippocampus, albeit its spread in the neocortex was limited. It remains to be clarified why enlargement is restricted in the neocortex, and whether it may occur after repeated reactivation *in vivo*. In summary, spine structural plasticity occurs in a stringent manner in the neocortex *in vivo*, which may provide a cellular basis for slow learning in the cortex (Lisman et al., 2001).

## Materials and Methods

### Surgery for the *in vivo* mouse experiment

All animal procedures were approved by the Animal Experiment Committee of the University of Tokyo (Tokyo, Japan). Procedures were conducted in accordance with the University of Tokyo Animal Care and Use Guidelines. The surgical procedure was previously described (Noguchi et al., 2011). In brief, we anesthetized adult mice expressing GFP or YFP in a subset of neurons: Thy1 GFP in the M line [GFP-M] or YFP in the H line [YFP-H]. Eighteen mice, aged 148 ± 129 days (expressed as the mean ± the SD), were used for the enlargement condition (YFP-H, 14 mice; GFP-M, 4 mouse). Eight mice, aged 70 ± 19 days, were used for the shrinkage condition (YFP-H, 7 mice; GFP-M, 1 mice). They were anesthetized with intraperitoneal injections of urethane and xylazine at 1.2 g/kg body weight and 7.5 mg/kg body weight, respectively, which were supplemented with the subcutaneous administration of the analgesic ketoprofen (20 mg/kg body weight). A steel plate with a recording chamber was attached to the skull by using cyanoacrylate glue so that the recording chamber was attached to the skull just above the visual cortex (3.0 mm posterior, 2.5 mm lateral to the bregma) (Paxinos & Franklin, 2001). The plate was then tightly fixed to the metal platform. We then removed the skull using a pair of forceps and a dental drill, which was fixed to a stereotaxic instrument (Narishige, Tokyo, Japan). The dura mater was carefully removed using fine forceps and a microhook to minimize any pressure applied to the brain surface. We then placed a semicircular glass coverslip to cover approximately one-half of the exposed brain surface (*Figure 1A*). The coverslip was fixed using dental acrylic (Fuji-Lute BC; GC Corp., Tokyo, Japan) or a stainless wire. The mice were supplied with humidified oxygen gas and warmed to 37°C with a heating pad (FST-HPS; Fine Science Tools Inc., North Vancouver, Canada).

### Two-photon *in vivo* imaging and uncaging

*In vivo* two-photon imaging and uncaging were conducted using an upright microscope (BX61WI; Olympus, Tokyo, Japan) equipped with a FV1000 laser scanning microscope system (Olympus) and a water-immersion objective lens (LUMPlanFI/IR 60× with a numerical aperture of 0.9; Olympus). The system included two mode-locked femtosecond-pulse titanium-sapphire lasers (MaiTai; Spectra Physics, Mountain View, CA, USA). The laser was set at 720 nm and used for uncaging (Matsuzaki et al., 2001). The other laser was set at 980 nm and used for imaging. Each light path was connected to the microscope via an independent scan head and acousto-optic modulator. For the 3D reconstruction of the dendrite images, 21–40 XY images separated by 0.5 μm were stacked by summing the fluorescence values at each pixel. 4-Methoxy-7-nitroindolinyl (MNI)-glutamate or 4-carboxymethoxy-5,7-dinitroindolinyl (CDNI)-glutamate was custom-synthesized by Nard institute Ltd. (Amagasaki, Japan) or purchased from Tocris Bioscience (Bristol, UK) was perfused in the recording chamber in artificial cerebral spinal fluid (ACSF).

### *In vivo* enlargement of dendritic spines

For the *in vivo* spine enlargement experiments, the cortical surface was first superfused with magnesium-free ACSF (ACSF w/o Mg^2+^) containing 125 mM NaCl, 2.5 mM KCl, 3 mM CaCl_2_, 0 mM MgCl_2_, 1.25 mM NaH_2_PO_4_ 26 mM NaHCO_3_, 20 mM glucose, and 10 μM tetrodotoxin (Nacalai, Kyoto, Japan). This solution was bubbled with 95% oxygen and 5% carbon dioxide for approximately 30 ± 15 min (expressed as the mean ± the SD; 20 dendrites). The bathing solution was then changed to ACSF w/o Mg^2+^ containing 20 mM MNI-glutamate or 10 mM CDNI-glutamate and 200 μM Trolox (Sigma-Aldrich, St. Louis, MO, USA), thereby enabling its diffusion into the cortical extracellular space approximately 15 min before the uncaging experiments. Two-photon uncaging was aimed at the tip of the spines, and repeated 60 times at 1 Hz. The power of the uncaging laser was typically set at 10 mW with an activation time of 0.6 ms. We expected that transient currents similar to miniature excitatory-postsynaptic currents were roughly elicited at this laser power; however, in this experiment we did not fine-tune the power along the cortical depth (Noguchi et al., 2011). For each experiment, 2–8 spines (average, 4.6 spines) were stimulated along a dendrite. We studied 52 spines/15 dendrites/14 mice with MNI-glutamate, and 22 spines/5 dendrites/4 mice with CDNI-glutamate. The success rate of enlargement was 25% and 13%, respectively. The solution was pooled in a small reservoir (2 mL) (*Figure 1A*). We constantly added pure water (after determining its flow rate empirically) to the reservoir to maintain the osmotic pressure of the solution at approximately 320 mOsm/kg. The solution was warmed at 37°C on the chamber by using circulating hot water (*Figure 1A*). All physiological experiments were conducted at 37°C.

### *In vivo* shrinkage of dendritic spines

For the spine shrinkage experiments, the cortical surface was superfused with ACSF containing 2 mM CaCl_2_ and 1 mM MgCl_2_. The bathing solution was then changed to ACSF that additionally contained 200 μM Trolox and a caged compound (i.e., 20 mM MNI-glutamate or 10 mM CDNI-glutamate). We studied 39 spines/16 dendrites/7 mice with MNI-glutamate, and 5 spines/2 dendrites/1 mice with CDNI-glutamate. The success rate of shrinkage was 36% and 25%, respectively. Repetitive stimulation was conducted at 1–2 Hz for 5–15 min with laser powers similar to those used for enlargement (~10 mW). As a control, the stimulation was also conducted in the presence of 50 mM D-2-amino-5-phosphonovaleric acid (APV), which is an NMDA receptor antagonist with MNI-glutamate.

### Analysis of the spine volume

Spine head volumes were estimated from the total fluorescence intensity by summing the fluorescence values of stacked images of the 3-D data, as previously reported using Image-J software (NIH, Bethesda, Maryland, USA)(Noguchi et al., 2011). When the image showed axon fibers overlapping with the target dendrite at different image depths, the spine head volume in the dendrite was calculated by partially summing the fluorescence values of sequential five Z-images by taking the moving average of the image stack along the Z-direction to avoid axonal fibers. A dendritic spine is near the diffraction limit of a two-photon microscope; therefore, the partially summed values (2 μm range in the Z-direction) should contain nearly the entire spine volume. Thus, the maximum value of the Z-moving average images allows good approximation of the total Z-summing of the stacked images.

Dendritic spines have spontaneous fluctuations in fluorescence because of spontaneous morphological changes, motility, or measurement errors. To determine spine volume fluctuations, we calculated the CV of the *in vivo* images before glutamate uncaging (14.7% ± 16.1% for 227 spines in the enlargement condition and 12.5% ± 7.9% for 196 spines in the shrinkage condition). We set the limit values of the fluctuation of the baseline as 2 CV (i.e., 30% for the enlargement data; 25% for the shrinkage data) and discarded the data when the fluctuation exceeded the limit value. Stimulated spines and neighboring spines with a prestimulation fluctuation over this limit (i.e., unstable spines) were discarded, as were the neighbors of the unstable stimulated spines. For the spine volume analysis, the average spine volume during 10– 30 min (i.e., enlargement) or 20–50 min (i.e., shrinkage) after the onset of the stimulation was calculated and are indicated as the difference from the baseline volume.

### Analysis of the spine neck length and the spine length

The 3-D spine neck length was measured manually using Image-J software (Noguchi et al., 2011). An XYZ stack image was resliced at z = 0.1 μm, and an XZ image of the spine neck was created. The intensity profile was measured along the neck in the XZ image. The half-maximum position of the spine and the parent dendrite was the edge of the spine and the dendrite, respectively. The spine neck length was calculated by subtracting the radiuses of the dendrite and the spine from the distance between peaks of the spine and the dendrite (*Figure 1B and Figure 2–figure supplement 2B*). For the analysis of the spine length before and after stimulation, the length between the tip of the spine and the edge of the dendrite was measured on the Z-stack images (*Figure 2– figure supplement 1D*).

### Statistical analysis

All data are presented as the mean ± standard error of the mean (SEM) (n indicates the number of spines), unless otherwise stated. Statistical tests of the spines were conducted using Excel-Statistics software (Social Survey Research Information Co. Ltd., Tokyo, Japan), as indicated. Differences from the baseline values were analyzed using the Wilcoxon signed-rank test (*Figure 1F; Figure 2E; and Figure 2–figure supplement 1A*). Differences between groups were analyzed using the Mann–Whitney rank sum test (*Figure 2–figure supplement 1B*). The Pearson’s product-moment correlation coefficient was calculated for the scatter plots (*Figure 1G; Figure 1–figure supplement 2A–D;* and *Figure 2–figure supplement 2A–2D*). The significance of a correlation coefficient was analyzed by using the *t*-test.

## Acknowledgements

We thank C. Maeda, M. Ogasawara, H. Ohno, K. Tamura, M. Nakajima, C. Matsubara and T. Sasaki for their technical assistance. We also thank N. Ichinohe for the helpful discussion and support. This work was supported by Grants-in-Aids for Science Research (S) (grant no. 26221011 to H.K.), Scientific Research (C) (grant numbers 18K06497 and 26430005 to J.N. and grant no. 2640290 to N.T.), Scientific Research on Innovative Areas (grant number 26111706 to J.N. and grant no. 16H06396 to S.Y.) and the Cooperative Research Program of “Network Joint Research Center for Materials and Devices” (grant no. 20171030 to J.N.), and World Premier International Research Center Initiative (awarded to HK) from the Ministry of Education, Culture, Sports, Science and Technology ([MEXT]; Tokyo, Japan), and Core Research of Evolutional Science and Technology ([CREST]; Tokyo, Japan; grant no. JPMJCR1652 to H.K.) from Japan Science and Technology Agency ([JST]; Tokyo, Japan), and the Strategic International Research Cooperative Program (SICP), Brain/MIND, and Strategic Research Program for Brain Sciences projects (grant no. 17dm0107120h0002) from the Agency for Medical Research and Development (awarded to H.K.).

## Competing Interests

The authors declare no competing interests.

## Author Contributions

J.N. and H.K. are co-corresponding authors and designed the study; J.N. conducted most imaging experiments; A.N., H.U., T.H., S.Y,. and N.T. assisted in some imaging experiments and in the data analysis; J.N. and H.K. wrote the manuscript. All authors contributed to the editing of the paper.

---

**Figure 1—figure supplement 1.**
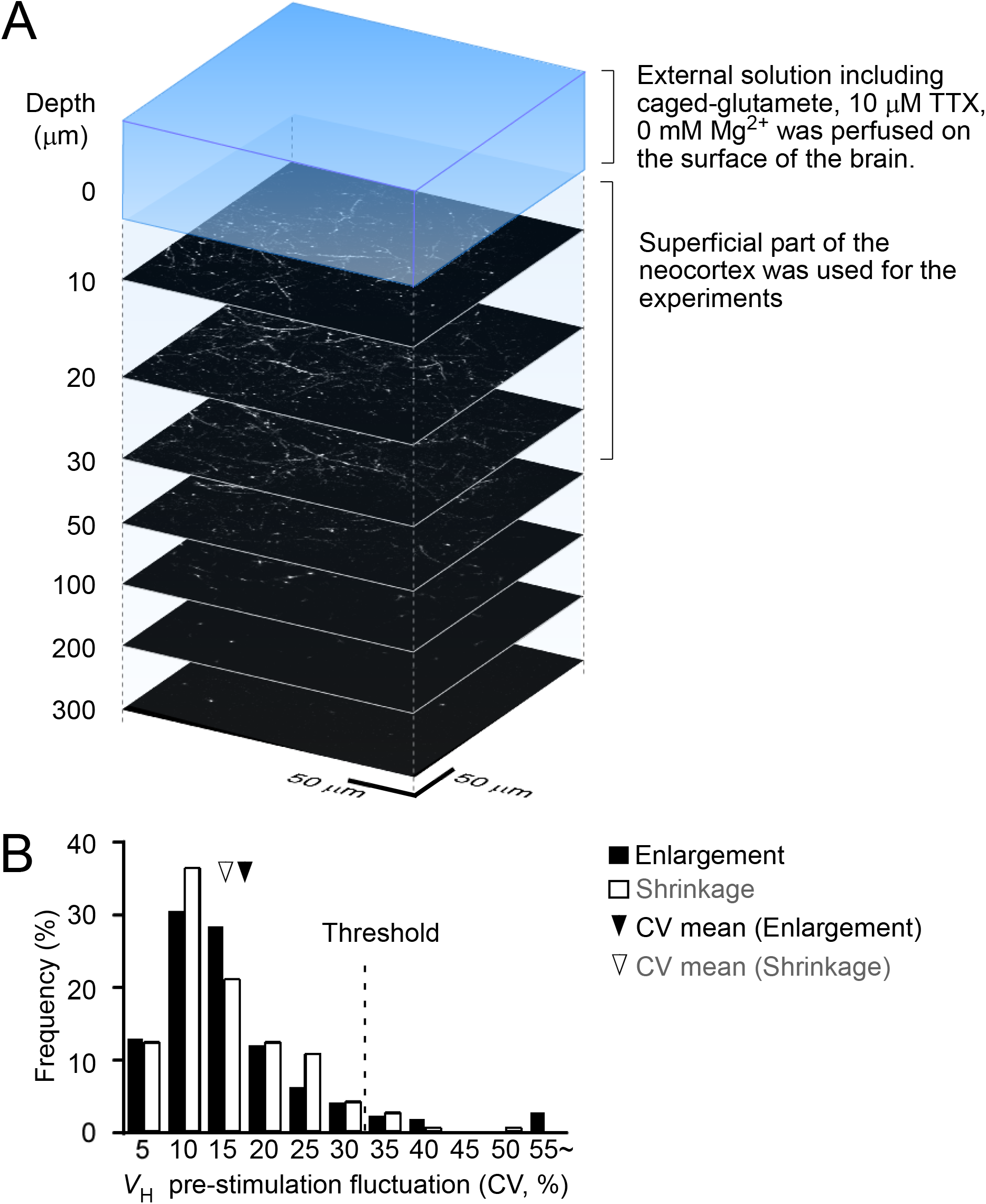
A diagram of the typical *in vivo* uncaging experiment. (**A**) The surface of the cortex is superfused with artificial cerebral spinal fluid (ACSF) solution containing 4-carboxymethoxy-5,7-dinitroindolinyl (CDNI) but devoid of magnesium (Mg^2+^). (**B**) The amplitude histogram of prestimulation spine volume fluctuations. The mean coefficient of variation (CV) is approximately 15% and is unaltered by the low Mg^2+^ solution. We thus set the threshold at 30% (i.e., 2 CV) for enlargement and for shrinkage.

**Figure 1—figure supplement 2.**
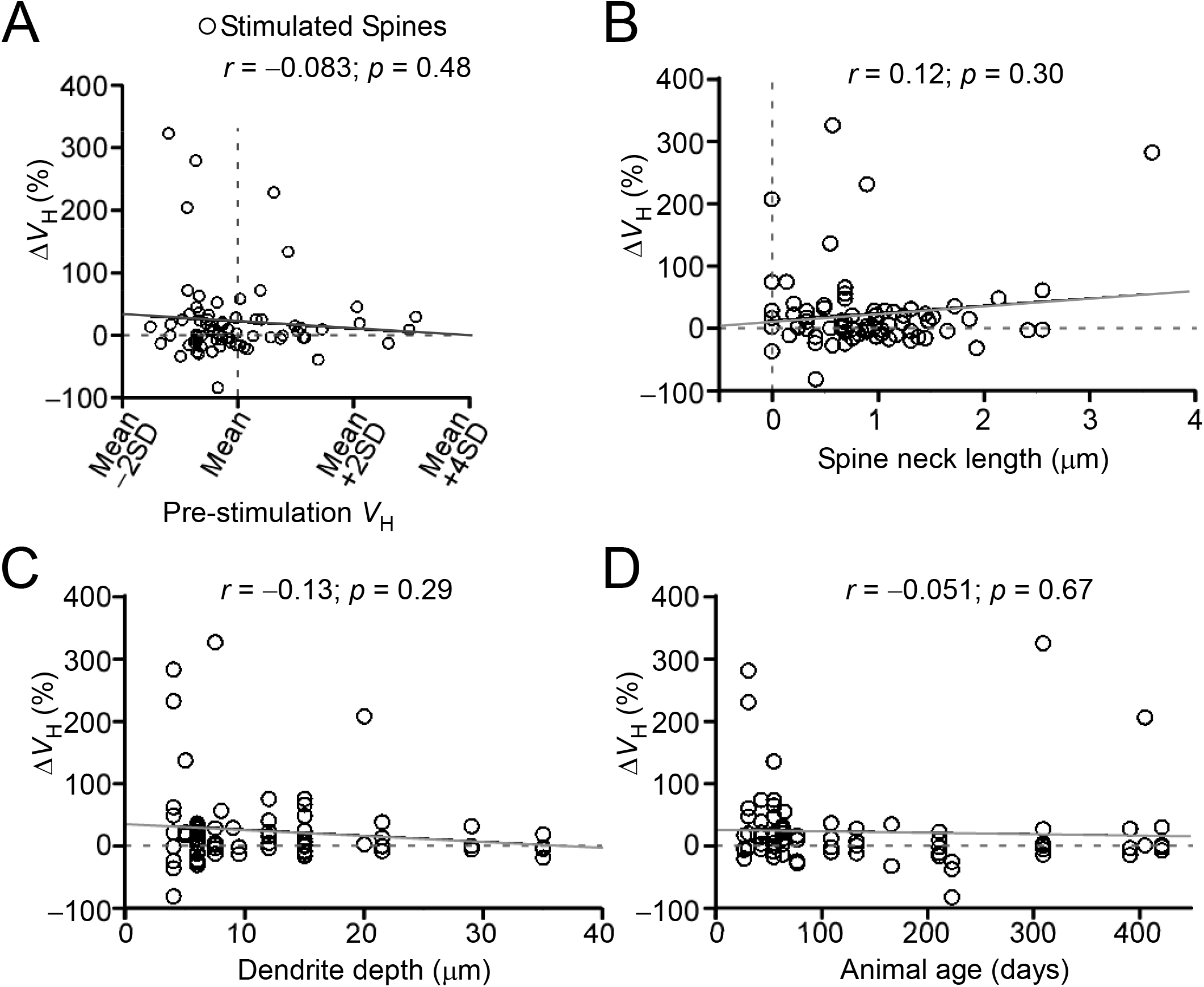
Investigation of the spine enlargement conditions. Scatter plots of the average spine enlargement of the stimulated spines (i.e., change in the head volume [ΔV_H_]) at 10–30 min after the stimulation against the relative prestimulation spine head volume for (A) each dendrite, (B) spine neck length, (C) dendrite depth, (D) and age of mice. Average enlargement of the stimulated spines from animals 0–60 days old (42.3% ± 14.2%, 26 spines, 5 mice), 61–200 days old (8.9% ± 4.7%, 23 spines, 6 mice), and 200+ days old (16.6% ± 16.0%, 25 spines, 7 mice). Pearson’s product-moment coefficient was calculated. The gray dotted and solid lines show the linear regression slopes.

**Figure 2–figure supplement 1.**
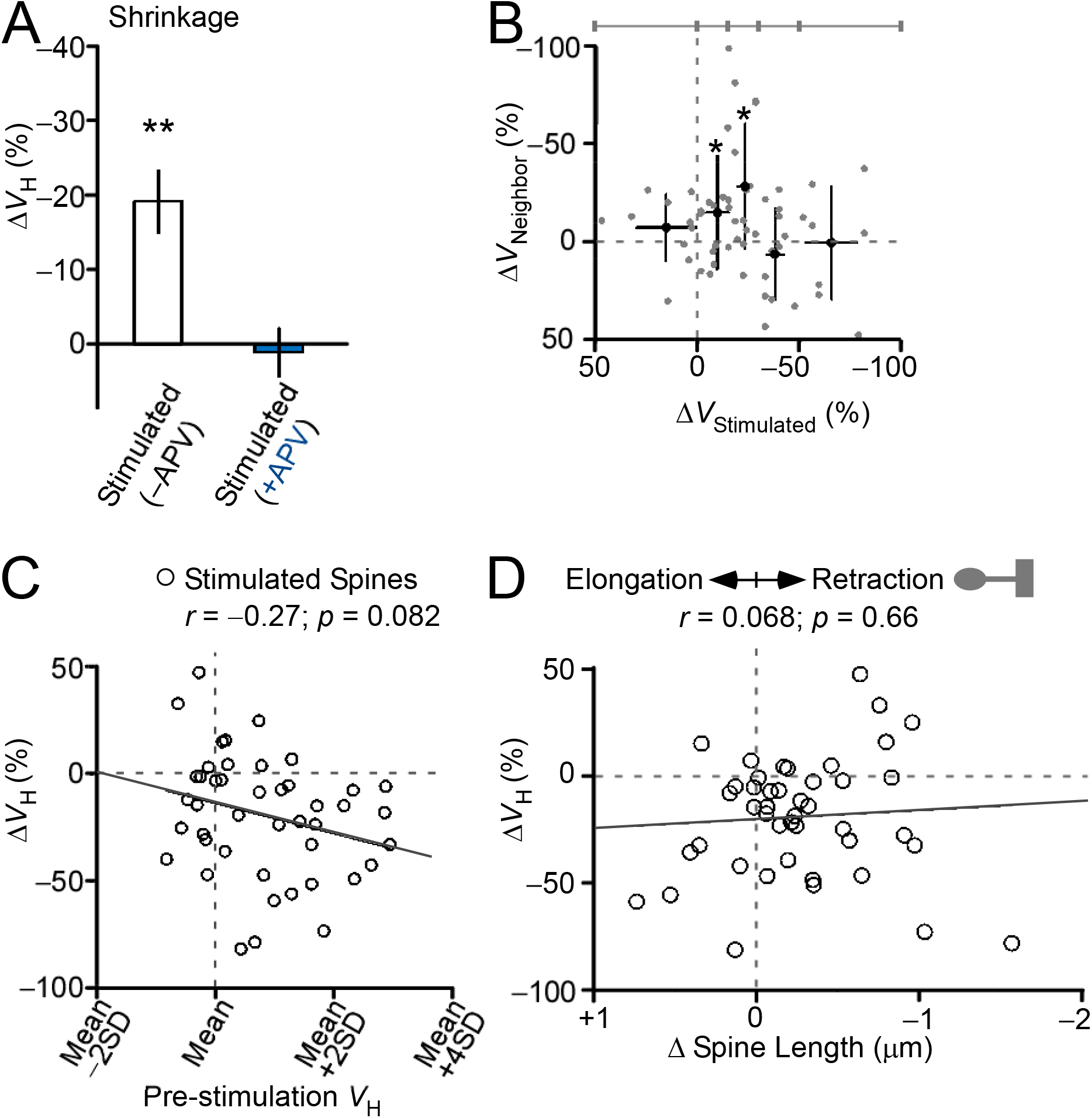
Properties of spine shrinkage. (**A**) The bar graph presents spine shrinkage (i.e., change in the volume head [ΔV_H_]) in the absence (19% ± 4%, 44 spines) and in the presence (1.1% ± 3.3%, 12 spines) of the NMDA receptor antagonist APV. The error bars represent the standard error of the mean (SEM). \*\**p* < 0.0001, based on Wilcoxon signed-rank test against zero. (**B**) The spine shrinkage of neighboring spines ([ΔV_Neighbors_]) at <3 μm is plotted against the spine shrinkage of the stimulated spines (ΔV_Stimulated_). The average values are calculated within the ranges of ΔV_Stimulated_, as indicated above the plot. The error bars represent the standard deviation (SD). \**p* < 0.05, based on Wilcoxon signed-rank test against zero. The scatter plots of the average spine shrinkage (ΔV_H_) of the stimulated spines against the spine properties present (**C**) the relative prestimulation spine head volume in each dendrite and (**D**) the spine retraction just after the end of the stimulation. The average retraction was −0.27 ± 0.07 μm (44 spines) and was significant (*p* = 0.0004), based on Wilcoxon signed-rank test against zero. Spearman’s rank correlation coefficient was calculated. The solid lines are the linear regression slopes.

**Figure 2–figure supplement 2.**
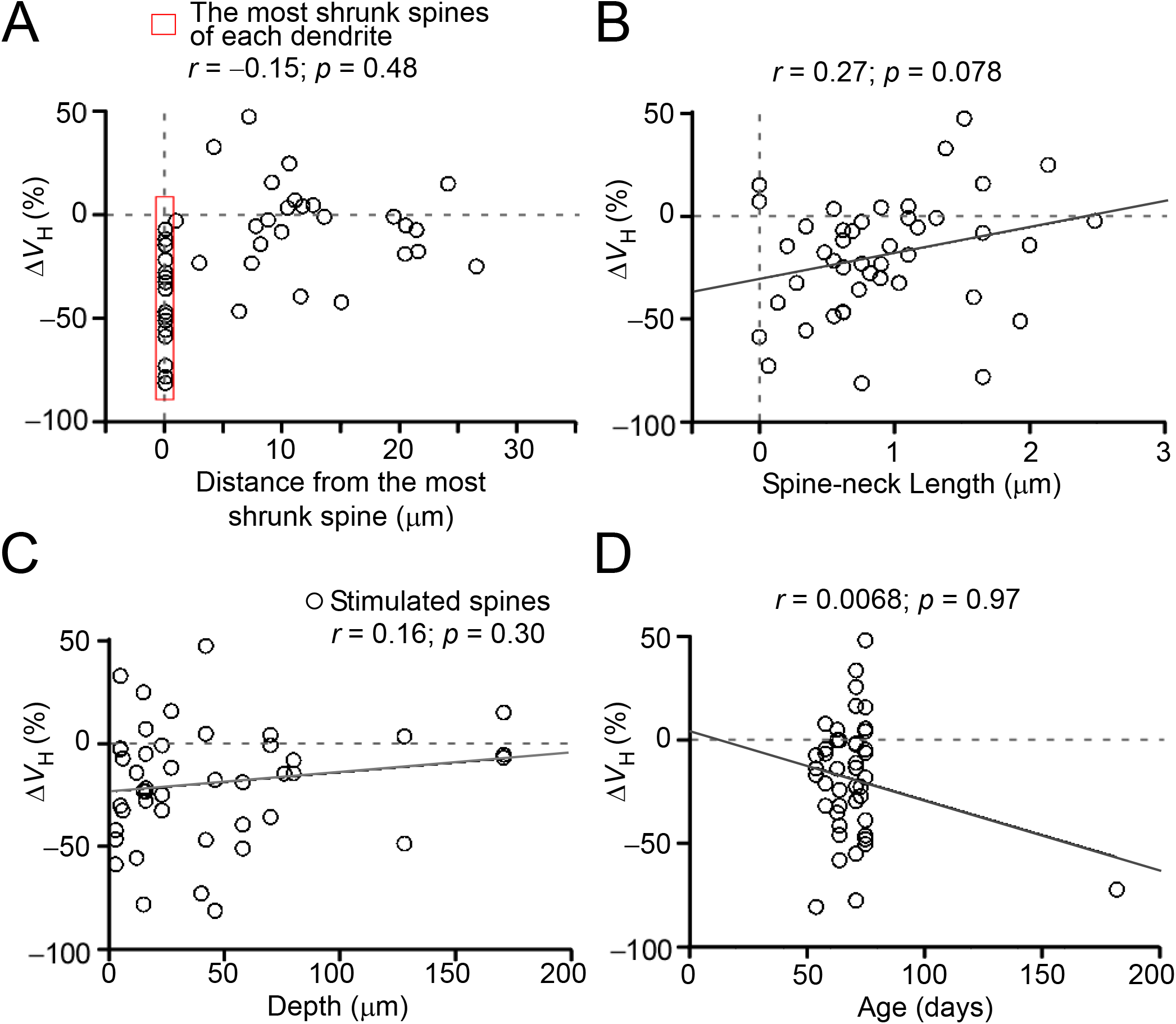
Conditions of spine shrinkage. Scatter plots of the average spine shrinkage (i.e., change in the head volume [ΔV_H_]) of the stimulated spines against (**A**) the distance between the most shrunken spine of each dendrite and other stimulated spines, (**B**) the prestimulation spine neck length, (**C**) the dendrite depth, and (**D**) the animal’s age (D). Pearson’s product-moment correlation coefficients were calculated. The solid lines present the linear regression slopes.

## References

Asrican, B., Lisman, J., & Otmakhov, N. (2007). Synaptic strength of individual spines correlates with bound Ca2+-calmodulin-dependent kinase II. J Neurosci, 27(51), 14007–14011. doi:10.1523/JNEUROSCI.3587-07.2007

Beique, J. C., Lin, D. T., Kang, M. G., Aizawa, H., Takamiya, K., & Huganir, R. L. (2006). Synapse-specific regulation of AMPA receptor function by PSD-95. Proc Natl Acad Sci USA, 103(51), 19535–19540. doi:10.1073/pnas.0608492103

Bhatt, D. H., Zhang, S., & Gan, W. B. (2009). Dendritic spine dynamics. Annu Rev Physiol, 71, 261–282. doi:10.1146/annurev.physiol.010908.163140

Bosch, M., Castro, J., Saneyoshi, T., Matsuno, H., Sur, M., & Hayashi, Y. (2014). Structural and Molecular Remodeling of Dendritic Spine Substructures during Long-Term Potentiation. Neuron, 82(2), 444–459. doi:10.1016/j.neuron.2014.03.021

Fiala, J. C., Spacek, J., & Harris, K. M. (2002). Dendritic spine pathology: cause or consequence of neurological disorders? Brain Res.Rev., 39(1), 29–54.

Forrest, M. P., Parnell, E., & Penzes, P. (2018). Dendritic structural plasticity and neuropsychiatric disease. Nat Rev Neurosci, 19(4), 215–234. doi:10.1038/nrn.2018.16

Govindarajan, A., Israely, I., Huang, S. Y., & Tonegawa, S. (2011). The dendritic branch is the preferred integrative unit for protein synthesis-dependent LTP. Neuron, 69(1), 132–146. doi:10.1016/j.neuron.2010.12.008

Harvey, C. D., & Svoboda, K. (2007). Locally dynamic synaptic learning rules in pyramidal neuron dendrites. Nature, 450(7173), 1195–1200. doi:10.1038/nature06416

Hayama, T., Noguchi, J., Watanabe, S., Takahashi, N., Hayashi-Takagi, A., Ellis-Davies, G. C.,…Kasai, H. (2013). GABA promotes the competitive selection of dendritic spines by controlling local Ca2+ signaling. Nat Neurosci, 16(10), 1409–1416. doi:10.1038/nn.3496

Hayashi-Takagi, A., Yagishita, S., Nakamura, M., Shirai, F., Wu, Y. I., Loshbaugh, A. L.,…Kasai, H. (2015). Labelling and optical erasure of synaptic memory traces in the motor cortex. Nature, 525(7569), 333–338. doi:10.1038/nature15257

Holbro, N., Grunditz, A., & Oertner, T. G. (2009). Differential distribution of endoplasmic reticulum controls metabotropic signaling and plasticity at hippocampal synapses. Proc Natl Acad Sci U S A, 106(35), 15055–15060. doi:10.1073/pnas.0905110106

Holtmaat, A., & Svoboda, K. (2009). Experience-dependent structural synaptic plasticity in the mammalian brain. Nat Rev Neurosci, 10(9), 647–658. doi:10.1038/nrn2699

Honkura, N., Matsuzaki, M., Noguchi, J., Ellis-Davies, G. C., & Kasai, H. (2008). The subspine organization of actin fibers regulates the structure and plasticity of dendritic spines. Neuron, 57(5), 719–729. doi:10.1016/j.neuron.2008.01.013

Kasai, H., Fukuda, M., Watanabe, S., Hayashi-Takagi, A., & Noguchi, J. (2010). Structural dynamics of dendritic spines in memory and cognition. Trends Neurosci, 33(3), 121–129. doi:10.1016/j.tins.2010.01.001

Kopec, C. D., Real, E., Kessels, H. W., & Malinow, R. (2007). GluR1 links structural and functional plasticity at excitatory synapses. J Neurosci, 27(50), 13706–13718. doi:10.1523/jneurosci.3503-07.2007

Lee, S. J., Escobedo-Lozoya, Y., Szatmari, E. M., & Yasuda, R. (2009). Activation of CaMKII in single dendritic spines during long-term potentiation. Nature, 458(7236), 299–304.

Lisman, J., & Morris, R. G. (2001). Memory. Why is the cortex a slow learner? Nature, 411(6835), 248–249.

Matsuzaki, M., Ellis-Davies, G. C. R., Nemoto, T., Miyashita, Y., Iino, M., & Kasai, H. (2001). Dendritic spine geometry is critical for AMPA receptor expression in hippocampal CA1 pyramidal neurons. Nat.Neurosci., 4, 1086–1092.

Matsuzaki, M., Honkura, N., Ellis-Davies, G. C., & Kasai, H. (2004). Structural basis of long-term potentiation in single dendritic spines. Nature, 429(6993), 761–766. doi:10.1038/nature02617

Noguchi, J., Hayama, T., Watanabe, S., Ucar, H., Yagishita, S., Takahashi, N., & Kasai, H. (2016). State-dependent diffusion of actin-depolymerizing factor/cofilin underlies the enlargement and shrinkage of dendritic spines. Sci Rep, 6, 32897. doi:10.1038/srep32897

Noguchi, J., Matsuzaki, M., Ellis-Davies, G. C., & Kasai, H. (2005). Spine-neck geometry determines NMDA receptor-dependent Ca2+ signaling in dendrites. Neuron, 46(4), 609–622. doi:10.1016/j.neuron.2005.03.015

Noguchi, J., Nagaoka, A., Watanabe, S., Ellis-Davies, G. C., Kitamura, K., Kano, M.,…Kasai, H. (2011). In vivo two-photon uncaging of glutamate revealing the structure-function relationships of dendritic spines in the neocortex of adult mice. J Physiol, 589(Pt 10), 2447–2457. doi:10.1113/jphysiol.2011.207100

Oh, W. C., Hill, T. C., & Zito, K. (2013). Synapse-specific and size-dependent mechanisms of spine structural plasticity accompanying synaptic weakening. Proc Natl Acad Sci USA, 110(4), E305–312. doi:10.1073/pnas.1214705110

Smith, M. A., Ellis-Davies, G. C., & Magee, J. C. (2003). Mechanism of the distance-dependent scaling of Schaffer collateral synapses in rat CA1 pyramidal neurons. J Physiol, 548(Pt 1), 245–258. doi:10.1113/jphysiol.2002.036376

Tanaka, J., Horiike, Y., Matsuzaki, M., Miyazaki, T., Ellis-Davies, G. C. R., & Kasai, H. (2008). Protein synthesis and neurotrophin-dependent structural plasticity of single dendritic spines. Science, 319(5870), 1683–1687.

Xu, T., Yu, X., Perlik, A. J., Tobin, W. F., Zweig, J. A., Tennant, K.,…Zuo, Y. (2009). Rapid formation and selective stabilization of synapses for enduring motor memories. Nature, 462(7275), 915–919. doi:10.1038/nature08389

Yagishita, S., Hayashi-Takagi, A., Ellis-Davies, G. C., Urakubo, H., Ishii, S., & Kasai, H. (2014). A critical time window for dopamine actions on the structural plasticity of dendritic spines. Science, 345(6204), 1616–1620. doi:10.1126/science.1255514

Yasumatsu, N., Matsuzaki, M., Miyazaki, T., Noguchi, J., & Kasai, H. (2008). Principles of long-term dynamics of dendritic spines. J Neurosci, 28(50), 13592–13608. doi:10.1523/JNEUROSCI.0603-08.2008

Zhou, Q., Homma, K. J., & Poo, M. M. (2004). Shrinkage of dendritic spines associated with long-term depression of hippocampal synapses. Neuron, 44(5), 749–757. doi:10.1016/j.neuron.2004.11.011

Zito, K., Scheuss, V., Knott, G., Hill, T., & Svoboda, K. (2009). Rapid functional maturation of nascent dendritic spines. Neuron, 61(2), 247–258. doi:10.1016/j.neuron.2008.10.054

